# Estimating encounter rates as the first step of sexual selection in the lizard *Anolis sagrei*

**DOI:** 10.1101/164863

**Authors:** Ambika Kamath, Jonathan B. Losos

## Abstract

How individuals move through their environment dictates which other individuals they encounter, determining their social and reproductive interactions and the extent to which they experience sexual selection. Specifically, females rarely have the option of mating with all males in a population—they can only choose among the males they encounter. Further, quantifying phenotypic differences between the males that females encounter and those that sire females’ offspring lends insight into how social and reproductive interactions shape male phenotypes. We used an explicitly spatiotemporal Markov chain model to estimate the number of potential mates of *Anolis sagrei* lizards from their movement behavior, and used genetic paternity assignments to quantify sexual selection on males. Females frequently encountered and mated with multiple males, offering ample opportunity for female mate choice. Sexual selection favored males that were bigger and moved over larger areas, though the effect of body size cannot be disentangled from last-male precedence. Our approach corroborates some patterns of sexual selection previously hypothesized in anoles based on describing them as territorial, whereas other results, including female multiple mating itself, are at odds with territorial polygyny, offering insight into discrepancies in other taxa between behavioral and genetic descriptions of mating systems.

## Introduction

Sexual selection is a layered process, with animals sequentially having to overcome intrasexual competition, intersexual mating preferences, and, for males, post-copulatory competition and choice before achieving reproductive success [1, 2]. Decades of research have spawned a vast literature on each of these aspects of sexual selection. However, the very first step of the mating sequence—encountering potential mates—is rarely quantified. How individuals move across space through time directly influences the number and phenotypic distribution of potential mates they encounter [2–4]. Moreover, by bringing about encounters between potential mates as well as between potential competitors, individuals’ movement patterns set the stage for subsequent sexual selection through male-male competition and female choice [5, 6]. Documenting animals’ movement patterns, to understand how often and which members of the opposite sex are encountered by individuals, is thus fundamental to discovering the extent to which sexual selection can act in the wild. Concurrently, quantifying phenotypic differences between potential mates (individuals encountered) and actual mates (individuals whose offspring are borne), yields insight into the nature of selection imposed by social and reproductive interactions.

In particular, individuals’ movement patterns determine the potential for female mate choice to drive sexual selection [7, 8]. Female mate choice has been studied extensively, yielding vigorous debate surrounding the precise mechanisms by which it arises, acts, and is maintained across a range of taxa (reviewed in [9–11]). Common to all models of female choice, however, is the idea that females can choose among males. But whether and to what extent individual females in fact have such a choice, and therefore the extent to which female choice can drive sexual selection, depends in large part on how many males they encounter [3, 12, 13].

In studies of sexual selection, examinations of movement behavior are often restricted to considering how females sample among males in taxa where female mate choice is already acknowledged to be important. For example, searching behavior is often thought to be pertinent to sexual selection in species where females must visit and choose among males in leks or at fixed display sites (e.g. [8, 14]). However, similar measurements of movement behavior, and of encounters between potential mates, are equally relevant to understanding the opportunity for sexual selection in other animal species, including those where female choice is not usually considered a major selective pressure [13, 15].

Unexpected opportunities for female choice are often uncovered in species in which earlier behavioral descriptions of mating systems, based on movement patterns and social interactions, are found to be inconsistent with more recent genetic descriptions of mating patterns [16]. For example, most birds were widely regarded as monogamous prior to the advent of genetic tools that revealed frequent extrapair copulation [17]. Occasionally, these inconsistencies have prompted researchers to reexamine animal movement to reconcile behavioral and genetic descriptions of mating patterns (e.g. [18, 19]). For example, tracking the movement behavior of red deer revealed that females move long distances between harems unexpectedly often, demonstrating the possibility of female choice in a system where sexual selection was thought to be dominated by male-male competition [20]. In general, though, discrepancies between behavioral and genetic descriptions of mating systems remain common—consider how often species are described as “socially monogamous,” for example, but “genetically promiscuous.” These discrepancies imply that our descriptions of movement and social behaviors in many species remain incomplete or inaccurate, and we do not fully understand how sexual selection has shaped and is shaped by these behaviors.

In this paper, we develop an explicitly spatiotemporal approach to estimate encounters between potential mates from observations of the movement behavior of male and female *Anolis sagrei* lizards. Our first goal is to investigate if females encounter multiple males, which could offer females the possibility of mate choice. This possibility has previously been considered unlikely in most anoles, which have widely been described as having a territorial social system in which males defend an exclusive, fixed space that contain female territories, implying that while males may mate with multiple females, most females mate with just the single male in whose territory they reside.

This description of *Anolis* as territorial and polygynous persists despite genetic data revealing widespread female multiple mating (reviewed in [21]). Our second goal is to characterize sexual selection in this population by examining the predictors of male reproductive success at two levels. First we ask if the number of potential mates encountered by males is associated with their phenotype (the spatial extent of their movement and body size). Second, we test three hypotheses to understand the phenotypic differences between potential mates and actual mates. We first examine if females bear offspring sired by the males they encounter more often [22]. Then, we ask if males encountered later in the breeding season are more likely to sire offspring than males encountered earlier (“last-male precedence”; [23]). Finally, given widespread sexual selection in animals for larger males [24] as well as pronounced male-biased sexual size dimorphism in *A. sagrei*, we ask if females disproportionately bear offspring sired by larger males.

## Methods

### Field sampling and egg collection

*Anolis sagrei* is a low-perching arboreal lizard native to Cuba and the Bahamas that has been established in Florida for nearly a century [25, 26]. Lizards were captured, marked, and monitored to estimate their movement patterns in the University Gardens on the University of Florida campus in Gainesville, FL, from March 4, 2015 to May 25, 2015 between 0900 and 1800 hours. Sampling began at the start of the breeding season when lizard activity increased post-winter, and concluded at about the time when female *A. sagrei* began laying eggs (based on our 2014 observations of hatchlings appearing at the end of June, after an approximately month long incubation period [27]). We caught most lizards within a 7140 m^2^ area and marked captured individuals with unique bead tags [28], which allowed us to subsequently observe and identify individuals from a distance without disturbing them (in total, 4% of observations were of unmarked individuals). When captured, we measured each individual’s snout-vent length (SVL) as a measure of body size, and removed ~2-3 cm of tail tissue for genetic analysis. At each subsequent observation of a lizard, we noted its identity and the time of the observation. We avoided observing the same individual more frequently than once per hour, allowing ample time for lizards to resume normal behavior if disturbed by us. At each observation, we also recorded the lizard’s spatial location (usually a tree; in areas of continuous vegetation, locations > 1m apart were considered distinct). Locations at which lizards were seen were mapped by triangulation based on measuring distances between locations. We also mapped the locations of all trees within the site at which lizards were not observed; we could thus include all trees to which a lizard could potentially have moved in our estimations of movement patterns (Figure S1). Approximately once a month, we recaptured and re-measured males to estimate the average growth rate of males in this population.

At the end of the observation period, we captured 36 marked females and housed them singly under established anole husbandry conditions [29] until mid-November. Each cage contained a pot of soil in which the resident female laid eggs fertilized by sperm stored from her copulations in the field. Eggs were incubated for two to ten days, after which embryos were dissected out for genetic analysis.

### Movement pattern analysis

Analyses were carried out in R v. 3.3.2 [30]. We used a discrete-time Markov chain to model lizards’ movements between mapped locations. We divided daytime hours (0800 to 2000 hours; anoles are diurnal, so we assumed that the lizards did not move at night) over the sampling period (83 days) into 996 hour-long blocks. Observations were assigned to the bin closest to the time of the observation. Transition probabilities (*P*_*j*_) between locations *i* and *j* were modelled as exponentially declining with the distance between the locations (*d*_*ij*_), with rows of the transition matrix then normalized to sum to one (*N* is the total number of locations):

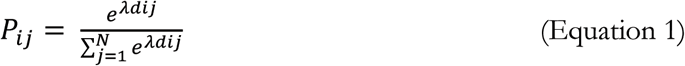

In other words, lizards were modelled as less likely to move to distant locations than to close locations, with a total probability of 1 of moving from each location to *some* location within the site, including staying at the same location. The value of the exponential decline parameter X was estimated by maximizing the likelihood of the observed data (including only pairs of consecutive observations of lizards, i.e. no assumptions were made while fitting the model regarding mortality or emigration after the last observation of a lizard) using the *bbmle* package [31]; separate models were fit for males and females.

Next, we used this Markov chain model describing the probabilities of lizards moving from one location to another to infer the probability that a lizard was at a particular location at a particular hour. Methodological details are provided in the Supplementary Information, but briefly, this probability depended on both where that lizard was seen previously and where it was seen next. We thus calculated, for each lizard, a matrix of probabilities that the lizard occupied a particular location at a particular hour, for all locations and hours. Rows of this matrix were normalized to one. Then, we performed element-wise multiplication of pairs of these matrices to calculate the probability of co-occurrence at each one hour-long time bin, for every possible pair of lizards. Encounters were categorized for each pair at each hour (“yes/no”) from these co-occurrence probabilities by setting cutoffs, i.e. pairs were classified as encountering one another if their co-occurrence probability was above the cutoff. We based cutoffs on the co-occurrence probabilities calculated for “observed encounters,” defined as pairs of lizards observed at the same location within an hour of one another. Cutoffs for classifying encounters between a pair of lizards depended on the connectedness of the locations at which these lizards were observed (i.e. the locations’ proximity to nearby locations; see Supplementary Information for details).

To quantify potential mating opportunities for each individual, we calculated the number of females encountered by each male and the number of males encountered by each female, as estimated by our model. The proportion of females that encounter multiple males and the mean number of males encountered by females reveal the extent to which multiple mating by females is possible in this population.

We quantified the spatial extent of an individual’s movement by calculating the mean of the distances from each observation of the individual to the centroid of all observations of the individual (mean distance from centroid). Lower mean distance from the centroid indicates smaller spatial extent. We jittered points randomly within a 0.5 m radius along both the X and Y axis of our site before calculating mean distance from the centroid, to account for the 1 m resolution at which locations were mapped.

We estimated a growth curve for males by fitting a logistic equation using nonlinear least squares regression [32] to males’ SVL measured initially and at recaptures, pooling data across all recaptured males (see Supplementary Information). We used this logistic growth curve to estimate the SVL of each male on the day of each of his inferred encounters, based on his SVL at the nearest capture, to test for sexual selection on male body size and for male avoidance of size-matched males (see below).

### Parentage analysis

DNA was extracted from the 36 females housed in captivity, all 161 sampled males, and 383 offspring using an AutogenPrep 965. Six microsatellite regions were amplified for these individuals (see Table S1 for primer and amplification protocol details; [33, 34]). Alleles were scored manually after examining chromatogram peaks in Geneious v10.0.9 [35].

Parentage analyses were performed in CERVUS v3.0.7 [36]. High proportions of null alleles were estimated at three loci (Table S1); following [37, 38], we retained these loci in the analysis but typed apparent homozygotes at only one allele, with the other allele coded as missing. All offspring had known mothers, and males estimated to have encountered the mother of a given offspring were considered candidate sires for that offspring. Further analyses (reported in the Supplementary Information) showed that simply restricting the number of candidate sires relative to the whole population did not inflate paternity assignments and that results of downstream analyses were unaffected by accounting for discordance between this analysis and an analysis where all males were provided as candidate fathers. In the simulation to determine log-likelihood ratio (LOD) cutoffs for paternity assignment, we provided a genotyping error rate of 0.01 (based on mother-offspring mismatch across all loci); the proportion of loci typed was 0.81. Simulations were run with 23 candidate sires and the proportion of sires typed set to 0.75, based on the maximum number of males encountered by any female in the population (17 males). These parameters were chosen after running preliminary analyses with simulations in which the proportion of sires typed was set to 0.25, 0.5, or 0.75 and the number of candidate sires set correspondingly to 68, 34, or 23; we chose the parameter combination with the closest match between the proportion of sires typed and the observed assignment rate [39].

### Hypothesis Testing

The number of potential mates encountered and spatial extent (mean distance to the centroid) had right-skewed distributions, and were log-transformed before parametric analyses; SVL was analyzed untransformed. T-tests and regressions were weighted by the number of observations per individual. We compared spatial extent between males and females using a t-test, and investigated if variation in males’ spatial extent was related to body size, using a linear regression of SVL at first capture vs. mean distance to the centroid.

Next, we examined if the number of females encountered by males varied with the spatial extent of males’ movement (mean distance from the centroid) and with mean male body size at their encounters with females, using a multiple linear regression.

To assess if males avoided size-matched males, we compared the differences in estimated SVL between pairs of males estimated to encounter one another vs. the differences in estimated SVL between randomly chosen pairs of males. For males that were estimated to encounter one another, we used the logistic growth curve to estimate their SVL on the day of the encounter. We initially sampled five random pairs per pair of males estimated to encounter one another, estimated their SVLs on the same day as the corresponding encounter, and eliminated random pairs in which either individual had an estimated SVL less than the minimum observed SVL. We then repeated this random sampling a total of 3000 times to assess if the number of male-male encounters among size-matched males (estimated SVL difference of 0-2 mm) was significantly lower than expected by chance.

We used a resampling approach to examine whether (1) males who sired individual females’ offspring encountered the mother significantly more often than males who encountered the same females but did not sire offspring (encounter rate hypothesis), (2) males who sired individual females’ offspring encountered those females later than males who encountered the same females but did not sire offspring (last-male precedence hypothesis) and (3) males who sired individual females’ offspring were bigger than males who encountered the same females but did not sire offspring (body size hypothesis).

We first calculated the difference between means of the number of encounters between male-female pairs for sires and non-sires across all offspring. We also calculated, for each male-female pair, the last hour at which the pair encountered one another and the maximum SVL estimated for the male across all encounters between the pair as an measure of male body size, and then calculated the difference between mean hour of last encounter and mean body size between sires and non-sires. We then recalculated these differences between means after randomly assigning each offspring a sire from the set of males that encountered the mother of that offspring. Random sire assignments were performed in two ways. To address the encounter rate hypothesis, we sampled uniformly from the list of males encountered by each mother. Then, to address the last-male precedence and body size hypotheses, we sampled in proportion to the rate at which each male encountered each mother. The former allowed us to test if sires encountered mothers more often than did non-sires, and the latter allowed us to test if later-encountered males and bigger males sired offspring more often than earlier-encountered males and smaller males, after accounting for variation across males in encounter rates. Each resampling was repeated 10000 times.

## Results

A total of 253 individuals (161 males, 92 females) were caught and marked during the sampling period, and were observed a total of 5629 times. The number of observations per individual ranged from one to 128; the median number of observations per individual was 11 for males and 15 for females. An example of two individuals’ locations through time is shown in Figure 1.

We used a Markov chain to model lizards’ movements between locations in the site, where transition probabilities were modelled as exponentially declining with the distance between locations. We estimated λ values of ‐0.78 for males and ‐1.27 for females (see Equation 1), indicating that males were more likely than females to move longer distances. Using this Markov chain model to estimate individuals’ movement patterns and thereafter the probabilities of their co-occurrence (see Figure 1 for an example), we calculated that females encountered 5.1 ± 3.7 males (mean ± S.D.) and males encountered 2.9 ± 3.0 females; 78% of females and 60% of males encountered multiple individuals of the opposite sex (Figure 2). Males encountered 4.5 ± 3.6 other males.

**Figure 1.**
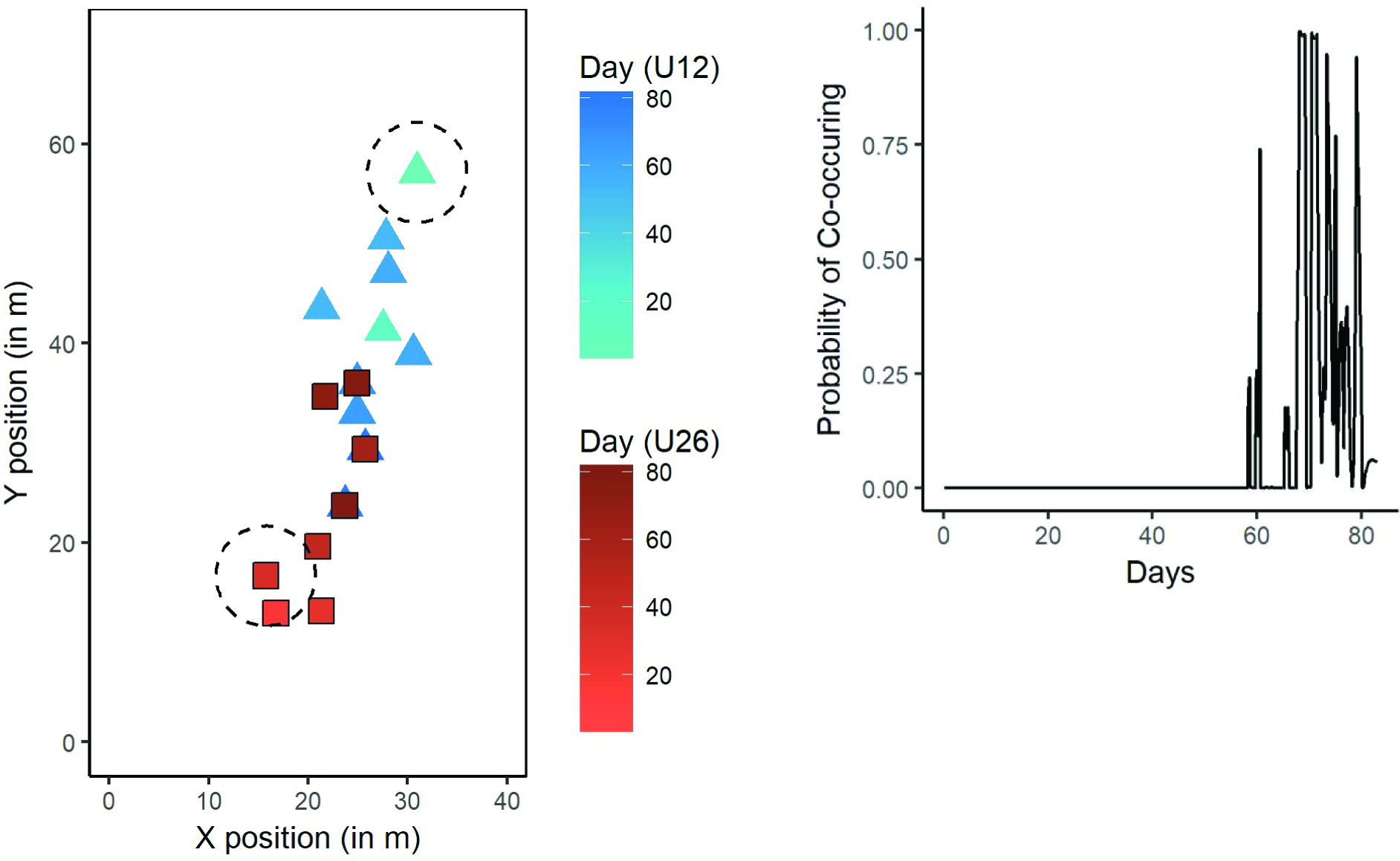
Examples of the locations through time for two individuals, U12 and U26 (left), and the probability of their co-occurrence as estimated by the Markov chain model (above). The dashed circles have a diameter of 10 m, which matches previous estimates of the territory size of male *Anolis sagrei*.

The mean distance from the centroid of all of an individual’s locations ranged from 0.2 m to 41.3 m for males and from 0.2 m to 20.8 m for females (mean ± standard deviation for males vs. females: 6.8 ± 7.0 m vs. 2.7 ± 3.3 m; *t* = 8.1, *df* = 208.7, *P* < 0.001). This measure of spatial extent was weakly associated with SVL at initial capture for males (r^2^ = 0.04, F_t,135_ = 4.91, *P* = 0.03).

**Figure 2.**
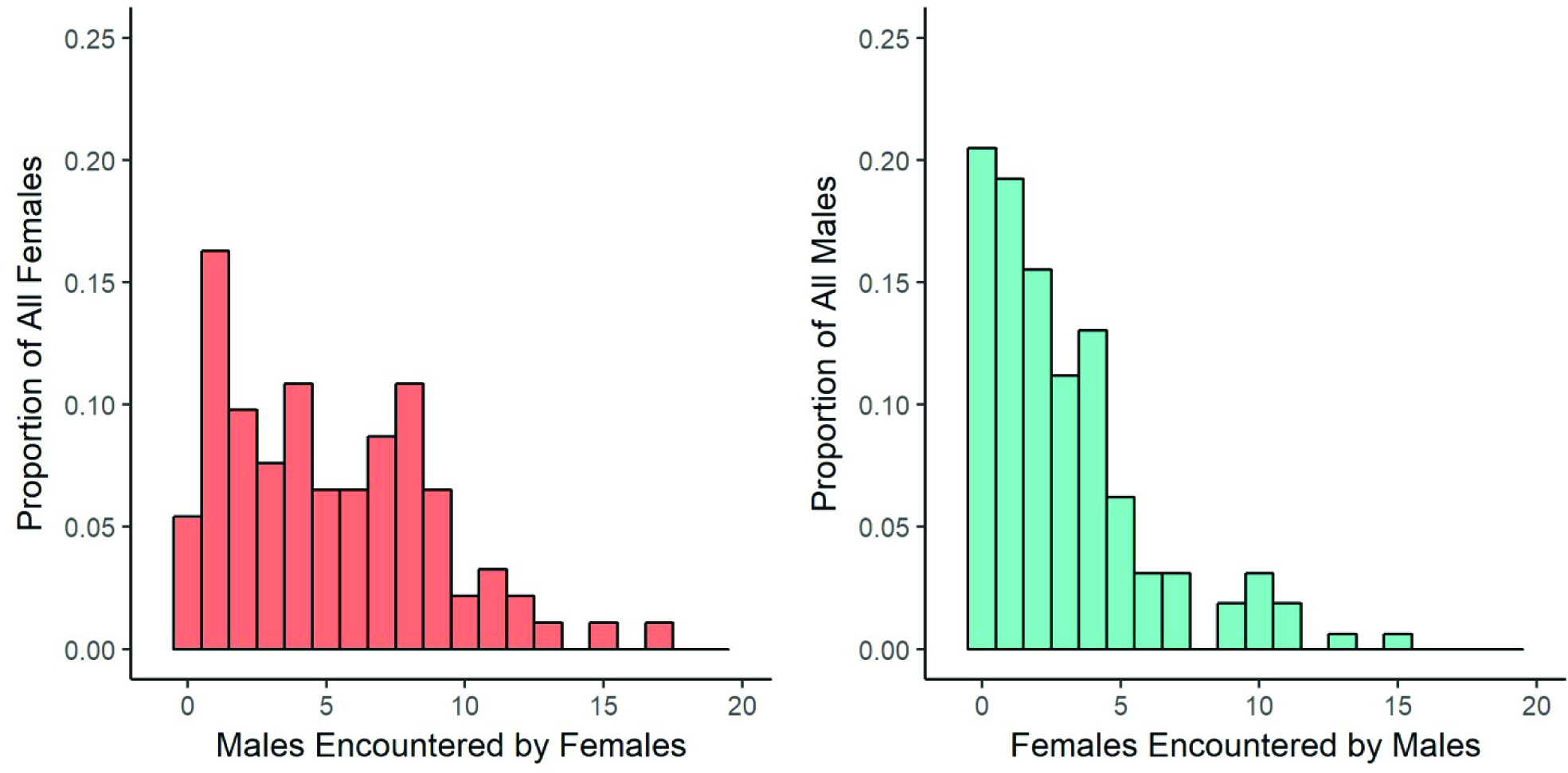
Histograms of the number of males encountered by females (left) and females encountered by males (right).

To estimate male body size at their encounters with females using a logistic growth curve, we recaptured 68 males and re-measured their SVLs a total of 94 times, with 32 ± 15 (mean ± SD) days elapsed between measurements. The mean difference in estimated SVL between pairs of males estimated to encounter one another (7.1 ± 4.5 mm) was comparable to the mean difference between randomly chosen pairs of males (7.4 ± 5.7 mm). However, observed size differences were underrepresented in the smallest (0-2 mm) category compared to random pairwise size differences (0.11 vs. 0.18 ± 0.002; *P* < 0.0003; Figure 3).

**Figure 3.**
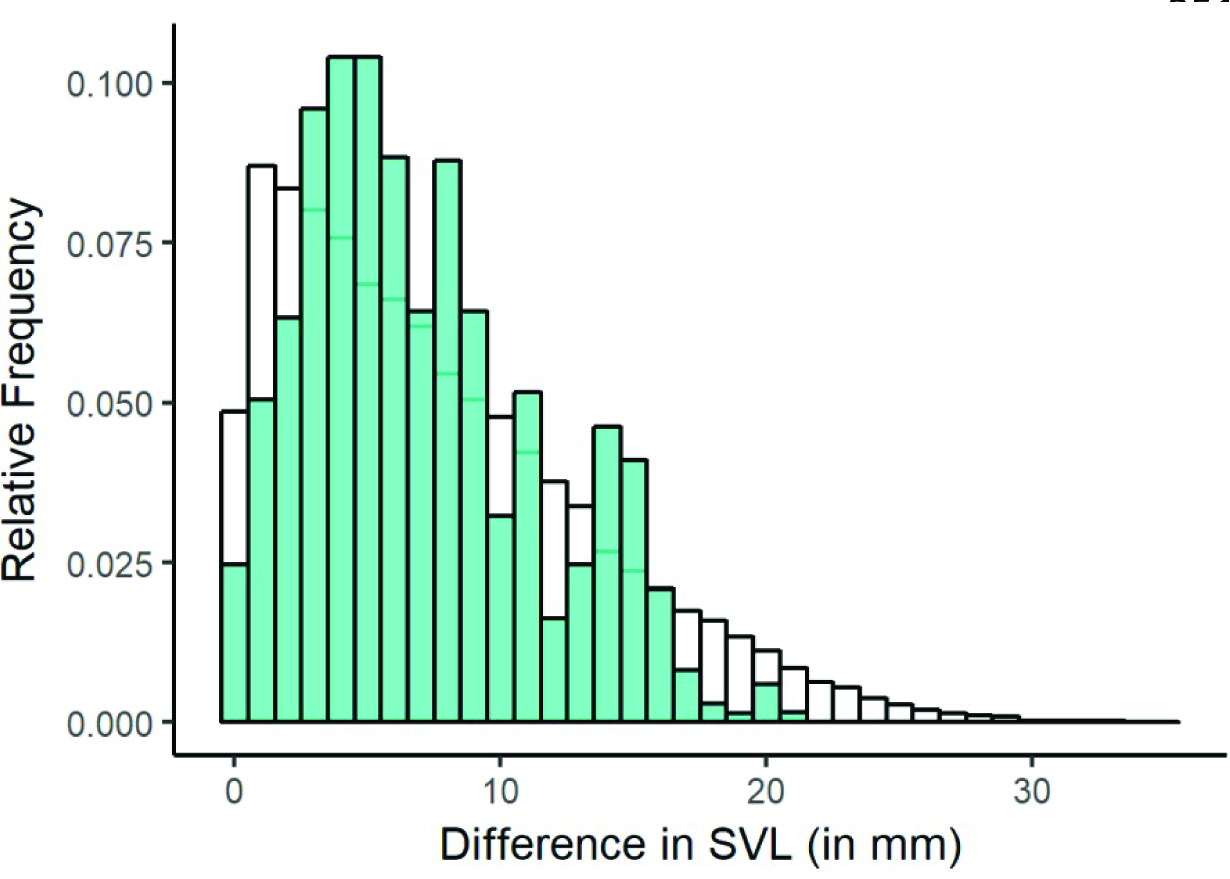
Estimated SVL differences between male pairs estimated to co-occur (blue bars), compared with pairwise SVL differences between randomly chosen pairs of males, with random males’ sizes estimated on the same days at the observed co-occurrences (white bars)

Males that encountered more females had a greater spatial extent (r^2^ = 0.10, F_1, 113_ = 13.0, *P* <0.001; Figure 4) and were larger in size on average at their encounters with females (r^2^ = 0.09, F_1, 113_ = 13.0, *P* <0.001; Figure 4); the interaction between spatial extent and SVL was not significant (r^2^ = 0.0, F_1, 113_ = 0.03, *P* = 0.87)

**Figure 4.**
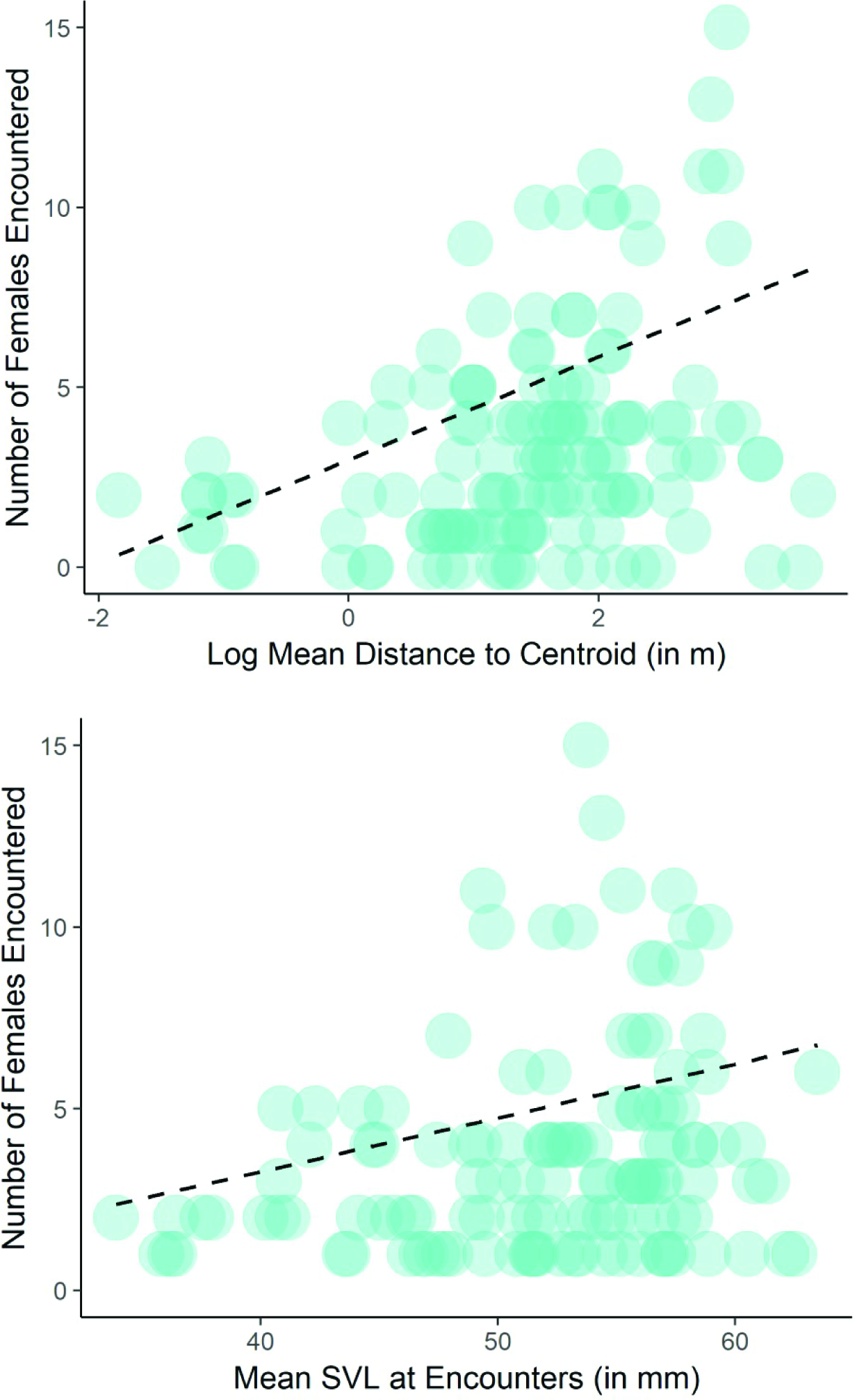
Relationship between the number of females encountered by males and the males’ spatial extent (top; measured as the mean distance to the centroid) and males’ mean estimated SVL across encounters (bottom).

Paternity was assigned to 84% of all offspring (323 individuals) at an 80% confidence level. We found that 64% of mothers bore offspring sired by more than one male; including offspring with unassigned sires, this proportion rose to 81%. Using a resampling approach to calculate p-values, we found that sires of offspring encountered mothers significantly more often than did non-sires (mean number of encounters between mothers and sires: 102 ± 140; non-sires: 40 ± 65; *P* < 0.0001). Accounting for variation across males in how often they encounter mothers, we found that sires encountered females significantly later than non-sires (mean ± SD of the last hour of encounter for sires: 892 ± 110; non-sires: 605 ± 258; *P* < 0.0001) and were significantly bigger than non-sires (mean ± SD of the maximum male SVL across encounters for male-female pairs, for sires: 57.8 ± 3.0 mm; non-sires: 53.2 ± 5.6 mm; *P* < 0.0001; Figure 5).

**Figure 5.**
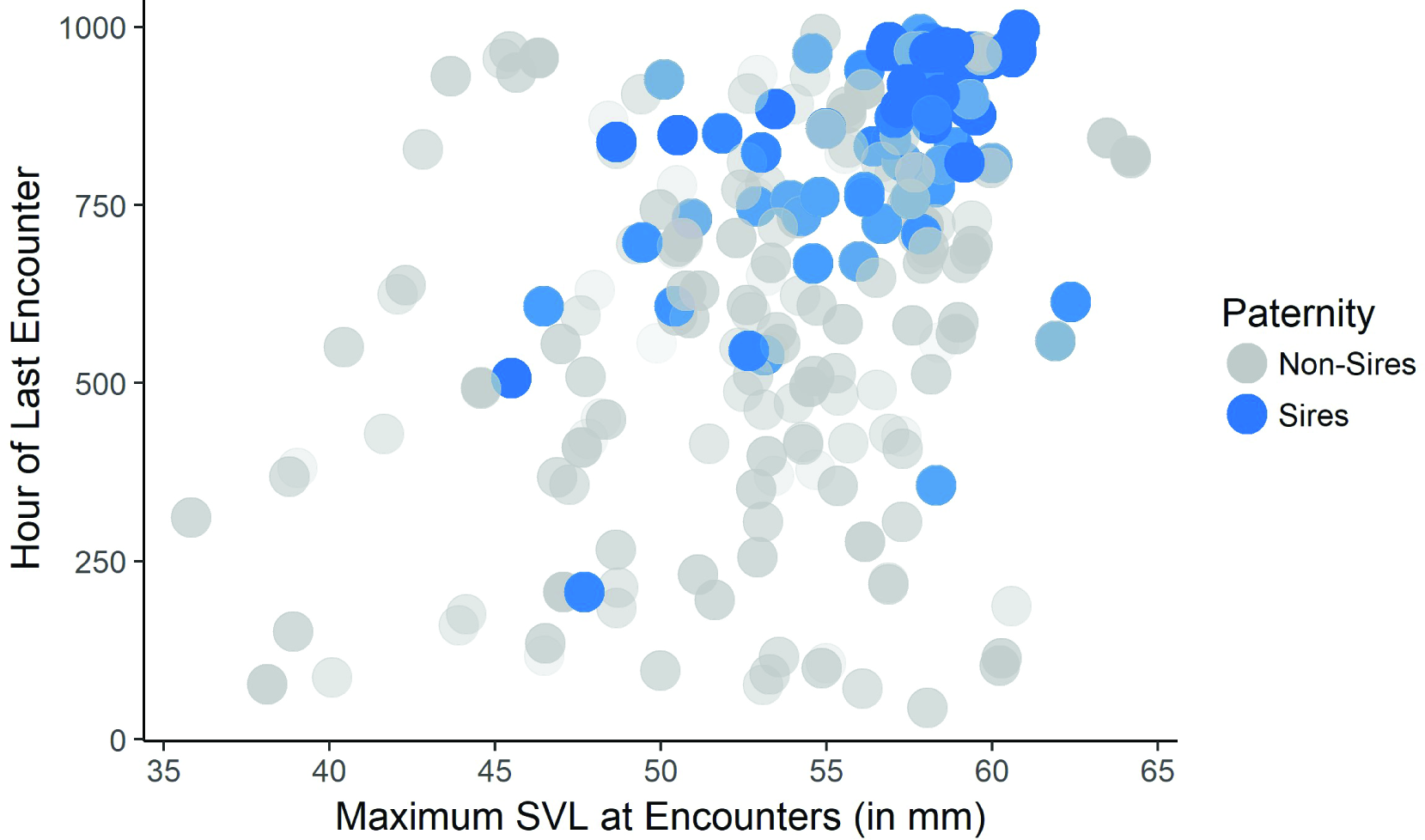
Maximum male snout-vent length (SVL) and the hour of last encounter for male-female pairs, colored by whether or not the male sired any of the female’s offspring.

## Discussion

How animals move through space determines how many and which other individuals they encounter, setting the stage for all subsequent social and reproductive interactions and ultimately determining reproductive success. Understanding animals’ movement patterns and the encounters they bring about is thus a key step in characterizing a population’s mating system, and is essential for determining how behavior both facilitates and is subject to sexual selection. Our spatiotemporal characterization of the movement patterns of a population of *A. sagrei* lizards revealed not only that a majority of males (60%) encountered multiple females but also that most females (78%) encountered multiple males over the first three months of the breeding season, indicating the potential in *A. sagrei* for complex polygynandrous mating patterns with ample scope for female choice.

Consistent with previous genetic descriptions of anole mating systems [40–42], we found that most females (64%–81%) bore offspring sired by more than one male. However, our results are at odds with most previous behavioral descriptions of movement patterns and mating systems in *Anolis* lizards, which leaned heavily on, and were constrained by, the framework of territoriality (reviewed in [21]). These behavioral descriptions were often coupled with an implicit expectation that anoles mate in a strictly polygynous manner, i.e., males mate with multiple females, but females mate with just the one male in whose territory they reside. Consequently, field studies have often implied that the opportunity for female choice in anoles is limited because it is precluded by territoriality (e.g. [43–45], but see [46]). Concurrently, despite varying evidence as to whether female choice is possible in natural populations, laboratory studies have offered females the choice between males to assess precopulatory mate preference [47–49] and have mated females with multiple males to assess postcopulatory sexual selection [50, 51]. Our results indicate that female anoles definitely have the opportunity to exercise post-copulatory mate choice and might also exercise pre-copulatory mate choice, calling into question the utility of territorial polygyny as a description of these lizards’ mating system.

Though rarely defined explicitly, territoriality is most often taken to mean the defense of an exclusive area in a fixed spatial location (reviewed in [52]). The inclusion of both exclusivity and site fidelity in the definition of territoriality is *necessary* for polygyny to be implied by territoriality. Departures from strict polygyny, as seen here and previously in anoles [40–42], imply departures from strict territoriality. While other definitions of territoriality, particularly those omitting site fidelity, could replace strict territoriality as a description of anole space use, these definitions do not imply strict polygyny, and indeed, make relatively few direct predictions about populations’ mating systems. We suggest that rather than trying to shoehorn descriptions of behavior into difficult-to-define concepts such as territoriality, we can reconcile widespread discrepancies between behavioral and genetic descriptions of mating systems by re-examining and quantifying animals’ movement and social behaviors as sequential steps in the process of sexual selection.

In territorial species, such re-examinations could further prompt us to revisit explanations of behaviors that have long been interpreted as characteristic of territorial polygyny, to discern if they might also be consistent with female choice. For example, we found that males encounter size-matched males less often than expected at random, which is consistent with male body size determining the outcome of male-male fights over territory ownership and access to mates [53, 54] and larger males subsequently excluding other large males from their territories [55, 56]. In this context of territoriality, smaller males are hypothesized to evade detection by larger territorial males, residing in their territories and attempting to ‘sneak’ copulations with resident females. But in taxa where females choose mates based on male body size (as is possible here; see below), larger males that retain smaller neighbors can accrue a mating advantage compared to males with neighbors of equal size (e.g. [57, 58]. Thus, males may engage in agonistic interactions to exclude size-matched but not smaller males from their vicinity, arranging themselves spatially relative to other males in a manner that raises their likelihood of being selected by females [59].

That said, our results support long-standing views about some facets of territoriality [52]. We found that males had a greater spatial extent than females, consistent with previous estimates of sex differences in territory size (e.g. [60, 61]), and with evidence for male-biased dispersal in anoles [62, 63]. But the spatial extent of individuals in this population is substantially higher than previous estimates of territory size in this and ecologically similar species (e.g. ~3 m^2^ for females and ~10 m^2^ to ~14 m^2^ for males [60, 64, 65], compared with mean minimum convex polygons areas of 36 m^2^ for females and 225 m^2^ for males in this study). Although it is possible that lizards in our site behaved unusually, the discrepancy may also partly be due to limited spatial and temporal sampling in previous studies, leading to underestimates of anole space use and interactions (reviewed in [21]). Indeed, subsampling from our dataset shows that if we had limited our spatial or temporal sampling extent to match the median sampling of previous studies, we would have detected a greatly reduced number of male-female pairs with overlapping home ranges (see Supplementary Information). In sum, we posit that while certain tenets of territoriality are well-supported in anoles, previous studies have likely underestimated the complexity of *Anolis* lizards’ movement patterns and social interactions by being constrained by a territorial framework. It remains unknown if similar problems afflict other species that have long been described as territorial.

Larger males not only encountered more females but were also more likely to sire offspring than smaller males, suggesting strong sexual selection for larger body size in male *A. sagrei*. These results are consistent with evidence across taxa that body size predicts male reproductive success [24]. However, most previous evidence in favor of this hypothesis in anoles is based on estimating mating patterns from home range area and overlap within the framework of territoriality (e.g. [60, 66, 67]. Our results show that the pattern of sexual selection favoring larger males is recovered even without a territorial interpretation of these lizards’ movement. As a species with largely indeterminate growth, body size can be an indicator of age [43, 68], or the ability to survive and thrive, suggesting an adaptive reason for females to choose to mate with, or bear offspring sired by, larger males [69]. However, selection on body size is difficult to disentangle from last-male precedence [23]—because males are smaller at earlier times in the breeding season, it is possible that large males sire more offspring simply because they have encountered and mated with females more recently.

Additionally, sexual selection may act on movement behavior—males with larger spatial extents encountered more females than males with smaller spatial extents. Because male body size and spatial extent were only very weakly correlated, the results do not indicate strong ontogenetic shifts in movement behavior. Thus, it appears that there are multiple ways for males to achieve reproductive success—they can grow large, they can traverse large areas, or both [70].

However, understanding the combined effects of body size, spatial extent, and last-male precedence is not necessarily straightforward—a single movement or mating strategy is unlikely to be adaptive in the face of social complexity. Instead, animals may make decisions about movement depending on their social and environmental context, rather than of adopting fixed patterns of space-use [5, 7, 8]. Such context-dependent decision making is often referred to as the maintenance of “alternative mating strategies,” though this variation need not be strictly discrete (e.g. [70, 71]). Individual-based models that incorporate the various sequential, compounding influences on reproductive success could reveal if males can make adaptive, context-dependent decisions to move or stay at particular locations based on their phenotype and the social and environmental situations they find themselves in.

Discerning if the decision rules used by individuals in a species are consistent across habitats and populations may represent a promising way of describing animal mating systems [72, 73]. In the future, sampling across populations and species that vary in density, sex ratio, and habitat structure will be essential to understanding how anoles’ decision rules regarding movement shape their mating system. The Markov chain approach to modelling movement patterns and estimating interactions presented here could be improved in such studies by incorporating both social and environmental variation influencing these decision rules—transition probabilities between locations could be scaled according to habitat preferences or the occupancy of a location by specific individuals, for example. At present, the accuracy of our estimation of encounters is limited by modeling movements as only based on distances between locations. Moreover, our approach is restricted to taxa that move relatively infrequently between relatively discrete locations, and cannot be readily modified for taxa with more continuous movement.

Finer spatial and temporal scales of location sampling, made possible with automated methods of tracking animals in the wild [74], could also allow for individual-level parameterization of any movement model, and would almost certainly improve the accuracy with which encounters are estimated. At present, a majority of the encounters (~70%) estimated by our model occur on the same day as a known observation of a lizard in the pair. This suggests that our model does not extrapolate unreasonably, but also shows that our discovery of these animals’ behavior remains limited by sampling. Ultimately, however, even finer scale location sampling will be insufficient for determining which encounters in fact lead to matings—discovering this crucial aspect of animals’ behavior will depend on focal animal observations of encounters in natural or naturalistic conditions. For a male, the gap between encountering a female and siring her offspring can include the sequential gauntlets of male-male competition, female mating preferences, and post-copulatory competition and choice [1, 2]. Disentangling effects on mating success at these various levels will be essential to fully understanding how sexual selection shapes behavior, and will require close observation of animal interactions prior to and including mating [75–77].

However, movement behavior is a precursor to bringing about any of these interactions—it comprises the first step of sexual selection and its quantification is therefore necessary for understanding the shape that sexual selection can take. In this paper, we develop a spatiotemporal framework to quantify movement behavior to estimate encounters between potential mates. This framework reveals an infrequently-recognized opportunity for female mate choice in *Anolis sagrei*, demonstrates that larger males are favored across multiple levels of sexual selection, and shows evidence for sexual selection on movement behavior and the timing of male-female encounters. We hope that similar frameworks centered on movement behavior can help to organize disparate studies that approach sexual selection at different levels in a variety of animal taxa.

## Acknowledgements

Jonathan Suh, Claire Dufour, Rachel Moon, Bonnie Kircher, and Barbara Tamires de Silviera assisted with data collection. Thom Sanger, Tom Wichman, Steve Johnson, and Stuart McDaniel provided logistical support. Thom Sanger additionally advised on lizard husbandry and embryo collection. Hanna Wegener, Jason Kolbe, Anthony Geneva, and Pavitra Muralidhar advised on the genetic methods and analyses. Ben de Bivort, Edward Burnell and Colin Donihue advised on the statistical analyses and, along with Yoel Stuart, Pavitra Muralidhar, Locke Rowe, Meera Lee Sethi, Cyndi Casey, and several anonymous reviewers, provided comments that improved the manuscript.

## Use of animals and field studies

This work was carried out under permit EXOT-15-09 from the U.S. Fish and Wildlife Service, and all procedures were approved under IACUC Protocol 26-11.

## Statement of Authorship

AK and JBL designed the study, AK collected data and performed analyses, AK and JBL wrote the manuscript.

## Data Accessibility Statement

data and code have been uploaded to Dryad (10.5061/dryad.kt387) and are currently available on GitHub (https://github.com/ambikamath/anolismovt).

## Supplementary Information

### Modelling lizard locations

As described in the main text, we modelled lizards’ movements between locations using a Markov chain model, where transition probabilities were fitted as exponentially declining with the distance between locations (Equation 1 in main text). If **M** is the transition matrix of the Markov chain model, and an individual lizard was observed at location A_m_ within hour T_m_ and then later at location A_n_ within hour T_n_, then we calculated the probability that that individual was found at a particular location (L) within the hour T ∈ (T_m_,…, T_n-1_) as follows:

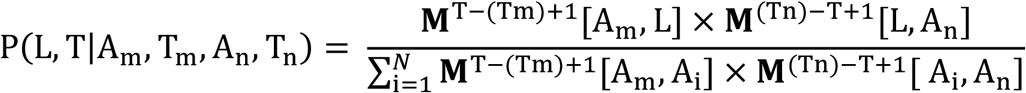

*N* is the total number of locations mapped; in our site, we mapped a total of 318 locations, which included not only locations at which lizards were observed but also trees within the site on which lizards were not observed but to which they might have moved (Figure S1). For the hours preceding the first observation and following the last observation for each individual, lizards were assumed to be equally likely to be anywhere in the site (i.e. the appropriate row or column of the transition matrix in equation 1 was replaced by a unit vector). This enabled us to include all sampled lizards, including those observed just once, into our estimation of encounters. It is worth nothing, though, that individuals with fewer observations contributed less to fitting the overall movement model than individuals with many observations. In sum, after this procedure, we obtained, for each lizard, a vector of probabilities of the lizard being in each of the 318 locations within each hour during the sampling period.

**Figure S1.**
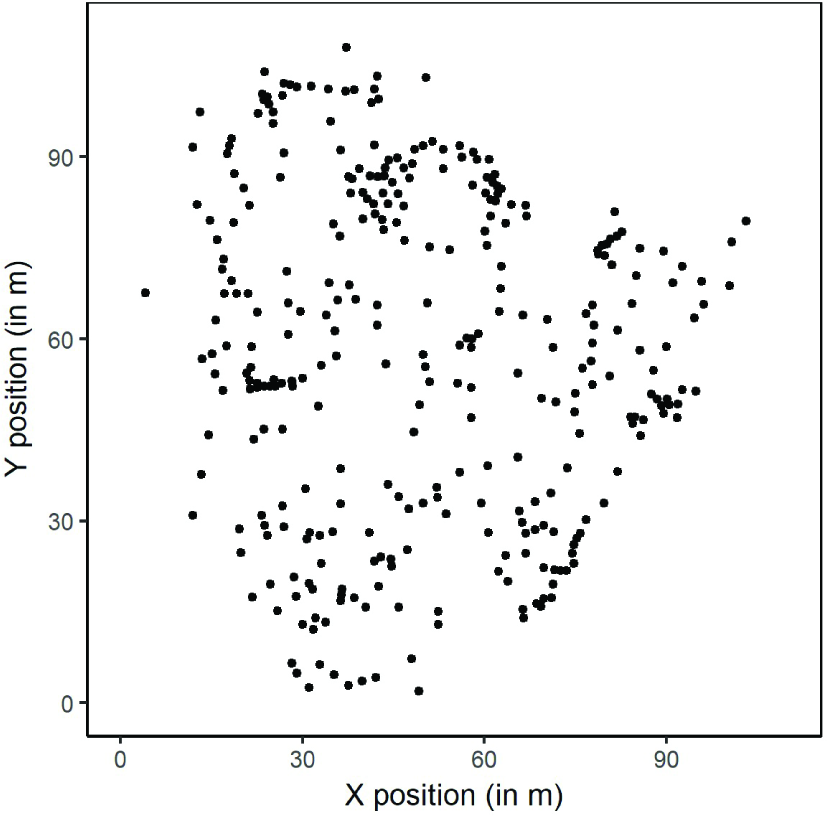
Map of all locations at which lizards were observed. In areas of continuous vegetation, locations more than 1m apart were considered distinct. We also mapped the location of all trees within the site at which lizards were not observed, to include all trees to which a lizard could potentially have moved in our estimations of their movement patterns, amounting to a total of 318 locations.

### Estimating Encounters

To estimate whether or not a pair of lizards encountered one another, we first performed an element-wise multiplication of the two matrices of the individuals’ probabilities of being at each location within each hour, and then summed probabilities across all locations for each hour in the resultant matrix for the pair. Probabilities were summed across locations so that two individuals that may be moderately likely to co-occur at multiple locations at the same time could still be estimated as encountering one another at that time.

We employed the following rationale and process to categorize encounters (“yes/no”) from co-occurrence probabilities:

- We first defined “observed encounters” as occurring between the pair of lizards observed in the same location as each other, within one hour of each other. We extracted, from the matrix described above, the co-occurrence probabilities of all observed encounters (n = 1028).
- Examining the variation in these co-occurrence probabilities of observed encounters, we noticed that they were related to the connectedness of the locations at which encounters took place, where connectedness was quantified as the mean distance from a location to the ten closest locations. Specifically, we saw that the calculated co-occurrence probabilities of observed encounters are lower for encounters that take place at more connected locations and higher for encounters at less connected locations (orange points in Figure S2). This is because when the probability of movement declines with distance, as modeled here, it is “easier” to leave a location that is close to other locations (i.e. more connected) than it is to leave a relatively isolated location (i.e. less connected). Consequently, the estimated probability of remaining at the former location will be lower compared to the probability of remaining at the latter location.
- We aimed to set co-occurrence probability cutoffs that would categorize all observed encounters as encounters, while also being as conservative as possible in converting co-occurrence probabilities into binary (“yes/no”) encounters. Balancing these two considerations, we set cutoffs for categorizing encounters on the basis of locations’ connectedness.
- Specifically, we binned locations on the basis of their connectedness (mean distance to the ten closest locations), into 1 m increments, and set distinct cutoffs for each bin, denoted by the black horizontal lines in Figure S2. In each connectedness bin, the cutoff for categorizing encounters in the remainder of the co-occurrence probability matrix was set to be the minimum probability calculated for observed co-occurrences in that bin. Had we used the global minimum of the calculated co-occurrence probabilities of observed encounters as a cutoff instead of the stepwise-increasing cutoffs described here (i.e. if the cutoff for the leftmost bin in Figure S2 were the cutoff for all co-occurrence probabilities regardless of location), we would have ignored the location-specific connectedness dynamics described above, leading us to infer many more encounters at less-connected locations.
- Each entry in the co-occurrence probability matrix was then assigned to a bin based on location connectedness. For all pairs of lizards for all hours in the sampling period, we considered the previous and next locations at which the lizards in the pair were observed, and found the minimum of the means of the ten closest distances for these locations. Depending on which bin this distance fell into, we assigned the appropriate probability cutoff for categorizing an encounter between that pair of lizards at that hour. If the calculated co-occurrence probability was higher than this assigned cutoff, the pair of lizards were categorized as encountering one another at this hour.

**Figure S2.**
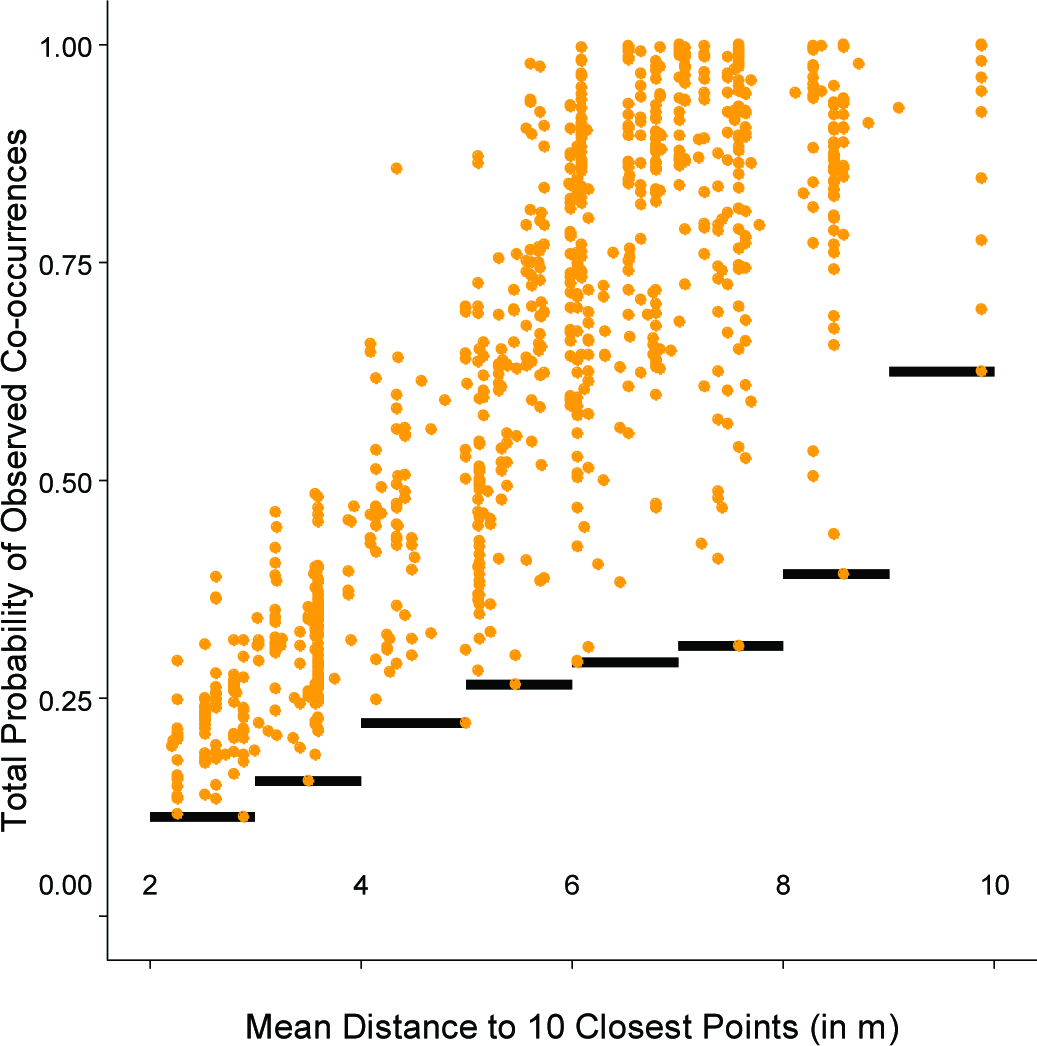
Cutoffs (black horizontal lines) for defining encounters, based on the mean distance from each location to the ten closest locations. Points indicate the sum of probabilities of co-occurrences across all locations, for pairs of individuals at hours where the two were observed at the same location within an hour of one another.

To assess the sensitivity of the number of male-female encounters to the probability cutoff described above, we repeated analyses using the 25% quartile instead of the minimum probability in each bin; a higher cutoff is more stringent, and will recover fewer encounters, but we were curious whether females would still be estimated as encountering multiple males across the sampling period. Not surprisingly, conservatively increasing the probability threshold for defining encounters reduced estimates of the number of potential mates encountered by both females (3.4 ± 3.1) and males (2.0 ± 2.1), but the potential for females to mate with multiple males remained.

Because boundary effects of a closed site might artificially increase the number of encounters, we also assessed if the number of male-female encounters was sensitive to treating the site as closed, by repeating the analyses described above with a simulated buffer of 50 locations, placed randomly within 20 m around the perimeter of the site. Allowing individuals to move to these locations did not alter estimates of the average number of males encountered by females (5.1 ± 3.7) or females encountered by males (2.9 ± 3.0).

### Growth Curve for Males

We estimated the growth curve of males in this population by fitting the following logistic equation (with parameters *a* and *r*) to the observed data, using nonlinear least squares regression (32Schoener and Schoener 1978):

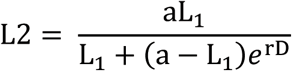

where L_1_ and L_2_ are the SVL of an individual measured at two successive captures, and D is the number of days between the two captures. Data from all recaptures of all males in the population were pooled to estimate this curve. We then used this logistic growth curve model to estimate the SVL of each male on the day of each of his encounters, based on his SVL at the nearest capture. Parameters for the logistic growth curve for male SVL (Equation 3) were estimated to be a = 63.7 and r = ‐0.016. Snout-vent lengths predicted by the model were highly correlated with measured SVLs (r^2^ = 0.92).

### Paternity analysis

Tests for sexual selection were performed on the paternity assignments in which candidate sires for offspring were the list of males their mother was estimated to have encountered (described in main text). However, we ran two further analyses. First, to assess if high paternity assignment rates in the main analysis were a consequence of restricting the number of candidate sires relative to the whole population, we also ran ten replicates of the analysis in which we sampled potential sires randomly from all 161 males, preserving the number of potential sires for each offspring individual and supplying the same group of potential sires for all offspring of a single female. Randomizing the identity but not the number of males encountered by each mother yielded 18% paternity assignment on average (13% to 24%), suggesting that the four-fold difference in assignment rates was due to identifying potential sires with greater accuracy based on their spatiotemporal movement patterns (assignment rate of 84% in the main analysis).

Second, we ran an analysis in which all sampled males were provided as candidate sires for all offspring. Simulation parameters specified for this analysis were 161 candidate sires and 0.96 proportion of males typed, based on the proportion of observations in the field that were of marked males. Paternity assignments were concordant for 80% of the individuals that were assigned paternity by both analyses. Of the remaining 39 individuals, the LOD scores for 21 mother-sire pairs differed between analyses by less than 1; these were considered false positives from the second analysis, and the assignments of the first paternity analysis were retained for downstream analysis. For the remaining 18 offspring, with mother-sire pairs where the LOD scores between the two analyses differed by more than 1, we found that results were not altered if we conducted downstream hypothesis testing both including and excluding their paternity assignments from the first analysis.

Twenty-six individuals (7% of the total) were assigned paternity only when all males were provided as candidate sires, i.e. their fathers were not estimated as having encountered their mothers. For these individuals, we calculated the minimum distance between any locations at which the mother and putative sire were observed, to assess the minimum distances that these individuals would have had to move to mate with one another. The minimum distance between locations at which mothers and sires of these 26 offspring were observed (regardless of *when* these lizards were found at those locations) ranged from 0 m to 56 m, with a median of 8.7 m.

We found no relationship (F_319_ = 0.52, P=0.47; Figure S3) between the day on which an egg was laid and sire SVL, suggesting that later laid eggs are not sired by poorer quality males, at least according to the one metric of male quality that we have data for, suggesting that females may not be compelled to use the sperm of less desirable males later in the laying season. We cannot rule out that sires of later laid eggs were worse in other ways than body size compared to sires of earlier laid eggs.

**Figure S3.**
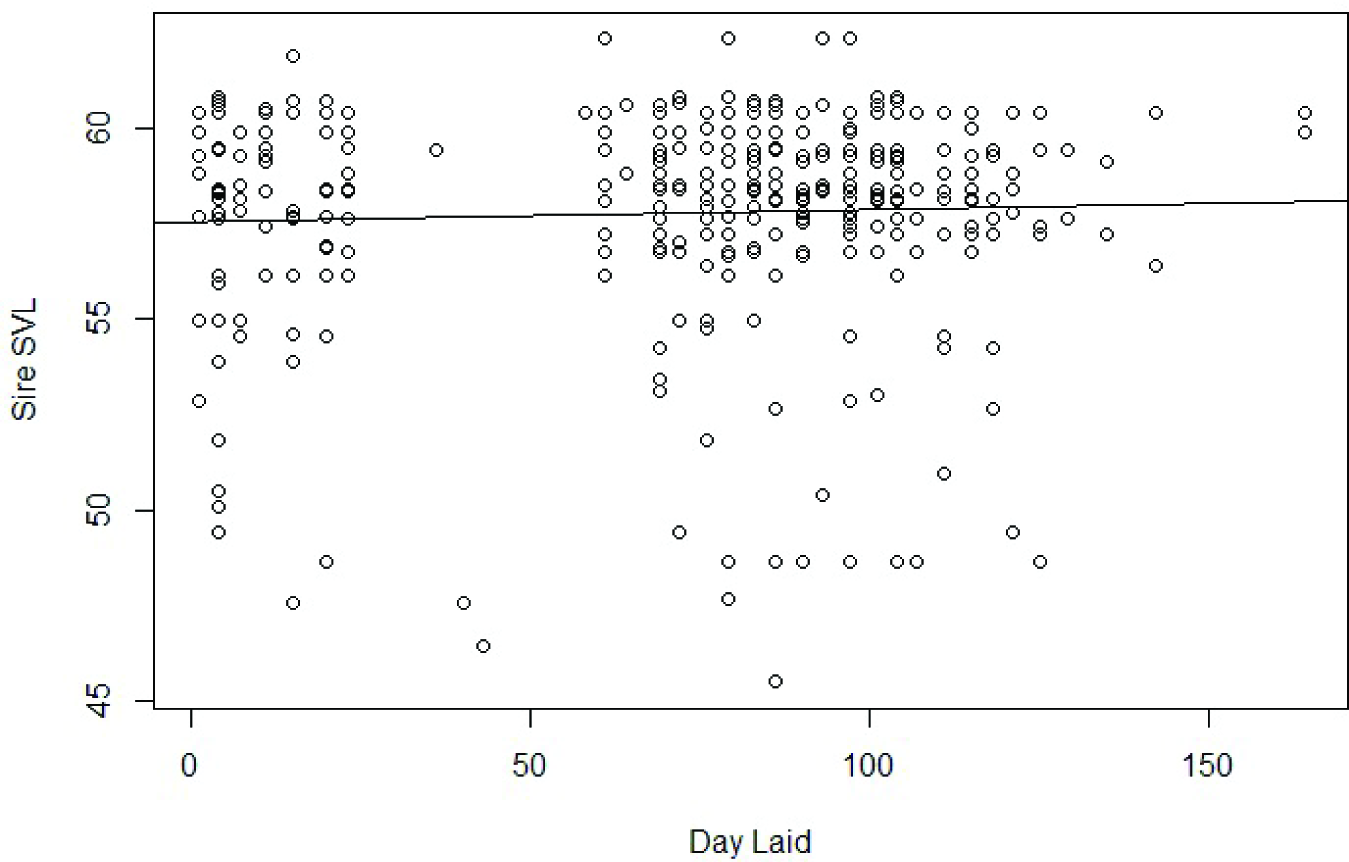
Relationship between the day on which an egg was laid and the SVL of the male siring that offspring, pooled across the 36 females from whom eggs were collected.

### Subsampling in Time and Space

To understand the effect of limiting sampling in space or time on male-female home range overlap (i.e. how mating patterns have often been assessed in previous studies of anoles), we asked if a purely spatial predictor— the overlap in minimum convex polygons (MCPs) between pairs of individuals—is affected by subsampling from our dataset to match either the median area (400 m^2^) or the median duration of sampling (4 weeks) or both of previous studies (see Table S2 for studies and sample areas and durations included in the calculation of the medians).

On average, in the whole dataset, female MCPs overlap with those of 12.8 ± 8.7 males, whereas male MCPs overlap with those of 8.1 ± 6.7 females. In subsamples of a randomly selected area of 400 m^2^ (repeated 404 times), we calculated that females overlapped a mean of 5.5 ± 3.1 males and males overlapped a mean of 2.3 ± 1.2 females. In subsamples of a duration of four weeks (for each of a possible 55 start dates), we calculated that females overlapped a mean of 4.5 ± 1.0 males and males overlapped a mean of 3.4 ± 0.8 females. In subsamples with a randomly selected area of 400 m^2^ and duration of four weeks, we calculated that females overlapped a mean of 2.4 ± 1.9 males and males overlapped a mean of 1.5 ± 1.0 females. Thus, limited sampling in space or time decreases the number of mates inferred from spatial overlap of MCP estimates of home range. That said, at all scales of subsampling, we recovered that females overlap with multiple males, possibly suggesting that our study population may be dissimilar to many of those studied previously, and also hinting that limited sampling in space and time is not the full explanation for why previous studies of *Anolis* territoriality have largely ignored the potential for females to encounter and mate with multiple males (but see Tokarz 1998).

**Table S1.**
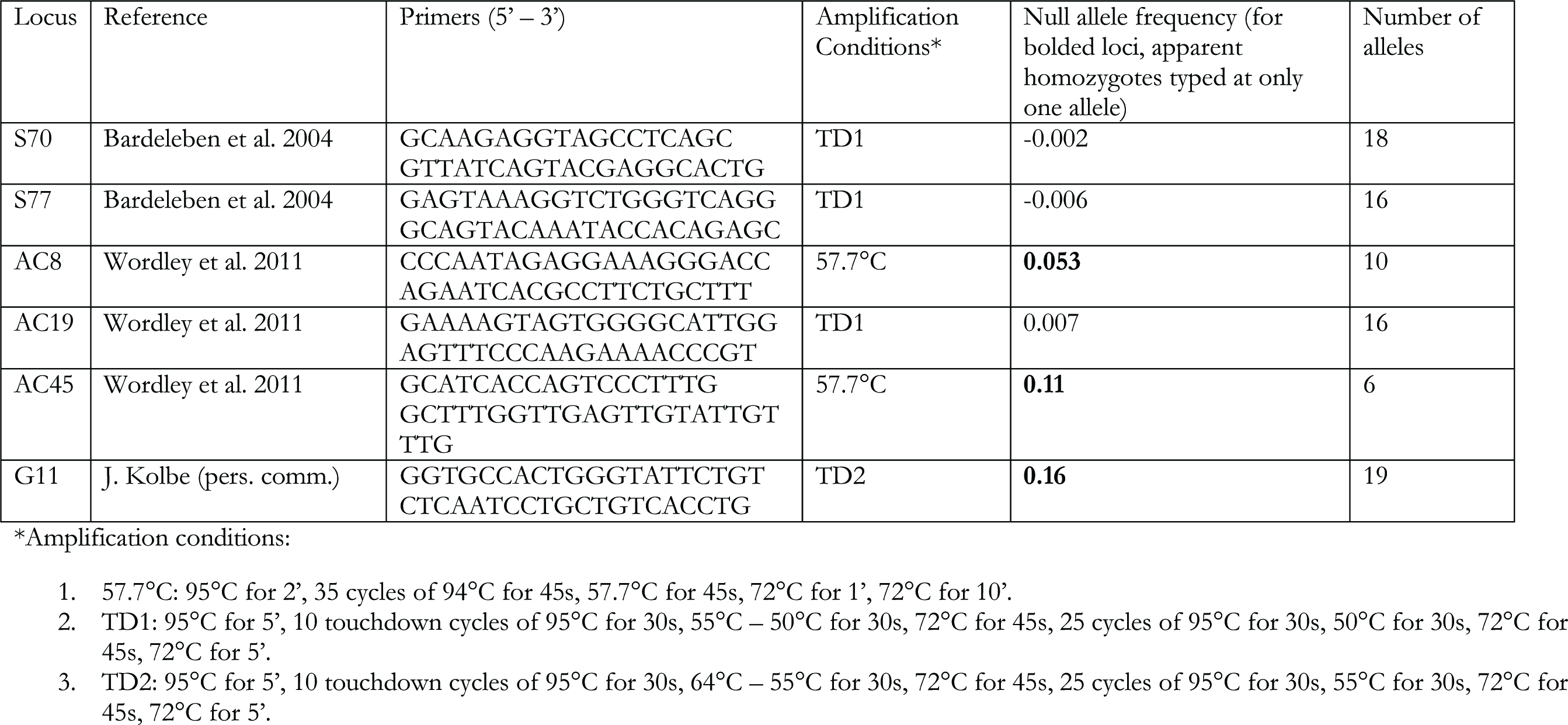

**Table S2.** Sampling area and duration of previous studies on *Anolis* home ranges Note that though the median duration is 3.5 weeks, a more conservative value of 4 weeks was used in the subsampling analysis above.

| Study | Sampling Area (in m<sup>2</sup>) | Sampling Duration (in weeks) |
| --- | --- | --- |
| Evans 1938 | 150000 | 3 |
| Greenberg and Noble 1944 | 25 | 47 |
| Gordon 1956 | 400 | 52 |
| Sexton et al. 1963 | 200 | 4 |
| Rand 1967 | 425 | 5 |
| Jenssen 1970 | 930 | 8 |
| Jackson 1973 | 14000 | 1 |
| Philibosian 1975 | 420 | 0.2 |
| Stamps 1977 | 9.5 | 5 |
|  | 34 | 13 |
|  | 47 | 13 |
|  | 37 | 15 |
| Hicks and Trivers 1983 | 12000 | 9 |
| Ruby 1984 | 441 | 28 |
| Fleishman 1988 | 169 | 10 |
| Jenssen and Nunez 1998 | 36 | 1 |
| Tokarz 1998 | 137 | 5 |
| Pereira et al. 2002 | 12 | 1 |
|  | 70 | 1 |
| Paterson 2002 | 400 | 1 |
| McMann and Paterson 2003 | 400 | 1 |
| Calsbeek 2009 | 1500 | 2 |
| Johnson et al. 2010 | 500 | 3 |
| Nicholson and Richards 2011 | 14000 | 52 |
| Bush et al. 2016 | 875 | 3 |
| Schoener and Schoener 1982 | 100 | — |
| Fitch and Henderson 1976 | — | 1 |
| Fitch and Henderson 1987 | — | 2 |
| Fobes et al. 1992. | — | 1.5 |
| **Median** | **400** | **3.5** |

